# *Wolbachia cifB* induces cytoplasmic incompatibility in the malaria mosquito

**DOI:** 10.1101/2021.04.20.440637

**Authors:** Kelsey L. Adams, Daniel G. Abernathy, Bailey C. Willett, Emily K. Selland, Maurice A. Itoe, Flaminia Catteruccia

**Affiliations:** Department of Immunology and Infectious Diseases, Harvard T.H. Chan School of Public Health, Boston, MA 02115

## Abstract

*Wolbachia* infections are a fascinating example of reproductive parasitism with strong potential to combat vector-borne diseases, due to their combined ability to spread in insect populations and block pathogen replication. Though the *Wolbachia* factors mediating the notable reproductive manipulation cytoplasmic incompatibility (CI) have now been identified as prophage WO genes *cifA* and *cifB*, the relative role of these genes is still intensely debated, with different models claiming that CI requires either both factors or *cifB* alone. Here we investigated whether *cifA* and *cifB* are sufficient to induce conditional sterility in the major malaria vector *Anopheles gambiae*, a species that appears to have limited susceptibility to invasion by *Wolbachia*. We report that CI can be fully recapitulated in these mosquitoes, and that *cifB* is sufficient to cause this reproductive manipulation. *cifB*-induced sterility is fully rescued by high levels of *cifA* expression in females. Surprisingly, however, when *cifA* is highly expressed in males alongside *cifB*, the CI phenotype is attenuated. *cifB* strongly impairs fertility also when expressed in the female germline, again mitigated by *cifA*. These data support a system whereby *cifB* and *cifA* must be fine-tuned to exercise CI and rescue, respectively, possibly explaining the limited success of *Wolbachia* at invading *Anopheles*. Our findings pave the way towards facilitating *Wolbachia* infections in anopheline vectors, for use in malaria control strategies.

## Introduction

*Wolbachia* endosymbionts are extremely successful insect colonizers due to reproductive manipulations they inflict on their insect hosts. One such notable manipulation is cytoplasmic incompatibility (CI), which is the failure of *Wolbachia*-infected males to produce viable progeny when mated to uninfected females^1^. Fertility is rescued in females colonized by *Wolbachia*, providing them with a reproductive advantage that, when paired with maternal transmission, allows these bacteria to effectively invade insect populations^2^. Recently the factors underscoring CI were identified as two genes, *cifA* and *cifB*, located adjacent to one another within WO prophage regions in the *Wolbachia* genome, with homologs in all known CI-inducing *Wolbachia* strains^3,4^. Since then, the activity and interactions between CifA and CifB have been disputed in the field. While it is generally recognized that *cifA* expression in the female rescues fertility, different theories (described in detail elsewhere^5–8^) hypothesize that either one (CifB) or both of these factors are necessary for inducing CI. The model whereby CifB is the toxin and CifA the antidote (generally referred to as the toxin-antidote model) is parsimonious as it implies only one role for CifA in both sexes, and is supported by *cifB* toxicity in yeast^4^. In *Drosophila melanogaster*, however, transgenic expression of *w*Mel *cifB* alone does not cause infertility, which is instead induced by co-expression of *cifA* and *cifB*. This finding lends support to a requirement for both genes in CI, known as the Two-By-One model^3,7^. While the toxin-antidote model can be modified to fit within a Two-By-One model framework^5,6^, its simplest interpretation does not, and there remains a lack of consensus on whether CifA is purely a rescue factor or a required accessory in CI.

Besides its role in reproductive parasitism, *Wolbachia* has attracted considerable attention for the control of vector-borne diseases due to the capacity of some strains to combine CI with strong pathogen-blocking effects. This is witnessed by large control programs based on the release of *Wolbachia*-infected mosquitoes to reduce transmission of dengue and other arboviruses by *Aedes* mosquitoes^9–11^. Additionally, the infertility induced by *Wolbachia*-infected males when mating with uninfected females can also be exploited to achieve suppression of insect populations, a strategy called Incompatible Insect Technique (IIT) that is also successfully applied in field trials of *Aedes* mosquitoes^12,13^. The implementation of similar *Wolbachia*-based programs to combat malaria transmission by *Anopheles* mosquitoes would be highly desirable, especially as widespread insecticide resistance threatens the resurgence of this devastating infectious disease ^14,15^. Unfortunately the utility of *Wolbachia* for malaria control is limited by a paucity of stable associations between *Anopheles* and these endosymbionts. Only one artificial *Wolbachia* infection has been achieved in the germline of an anopheline species, where however the *w*AlbB strain in *Anopheles stephensi* showed only partial rescue of CI and thus limited capacity for population invasion^16^. Furthermore, natural *Wolbachia* infections in field populations of important malaria vectors are reported sparsely and at low titers^17–22^ and there was no evidence for CI when investigated^23^. Such limited success at colonizing field and laboratory *Anopheles* populations has prevented the use of these bacteria for reducing the burden of malaria, yet the biological underpinnings of this phenomenon remain undetermined.

We investigated whether *cifA* and *cifB* are capable of inducing CI in *Anopheles gambiae*, the most important malaria vector in Africa. After codon-optimization, we separately cloned the two Type I *cif* genes from *w*Pip (also referred to as *cidA* and *cidB*^4^) under the *zero population growth (zpg)* promoter to drive expression in both male and female germlines^24^ (**Fig. 1a).** Co-injection of *zpg-cifA* and *zpg-cifB* constructs yielded F1 transgenics expressing either *cifA* alone, or both *cifA* and *cifB (zpg-cifA;B)*, but none that expressed *cifB* only, a suggestion that *cifB* may cause embryonic toxicity alleviated by *cifA* co-expression.

**Figure 1:**
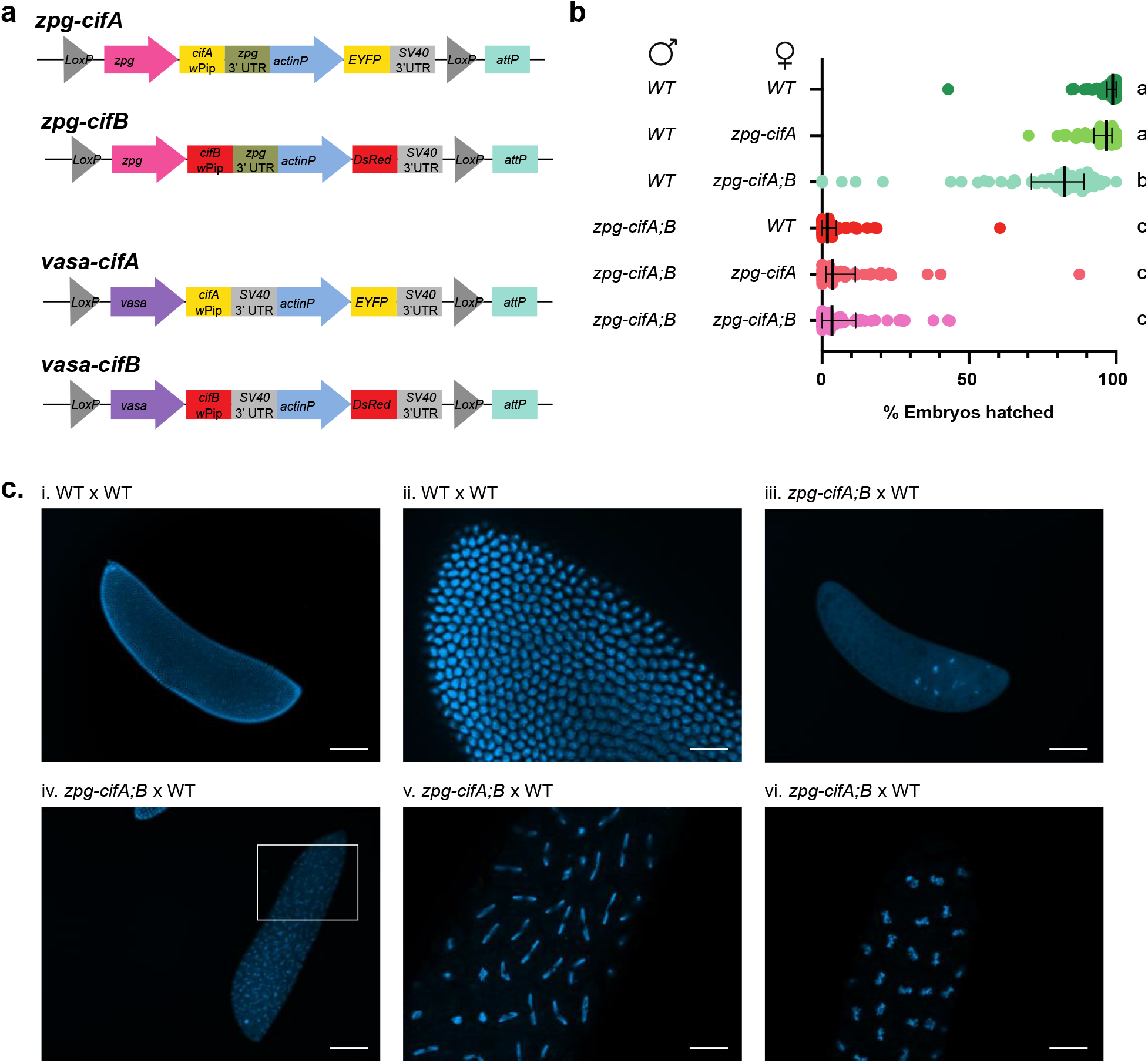
Co-expression of *cifA* and *cifB* in male *An. gambiae* causes embryonic lethality in progeny. (a) Construct design of *zpg-cifA, zpg-cifB, vasa-cifA*, and *vasa-cifB*. (b) Males that express *zpg-cifA;B* produce largely inviable progeny, regardless of whether their female mate expresses *zpg-cifA*. Expression of *zpg-cifA;B* in females causes a decrease in female fertility compared to WT, but *cifA* alone does not (Dunn’s multiple comparisons, *p*≤0.0071 for a vs b, *p*<0.0001 for a vs c and b vs c). Median and interquartile ranges are shown. For each group (top to bottom) the *n* (number of broods) is as follows: 58, 52, 59, 51, 53, 62. Kruskal-Wallis test: H=265, *p*<0.0001, df=5. (c) Embryos from *zpg-cifA;B* males crossed with WT females (or WT crosses, as controls in i. and ii.) were fixed and imaged with DAPI 3-4 hours post oviposition, showing developmental arrest of most CI embryos during early nuclear divisions (iii.), while some embryos complete multiple rounds of nuclear division but show mitotic defects such as chromatin bridging (iv. with a close-up in v.) and other chromosomal abnormalities resulting in delayed or arrested development (vi). Scale bars represent 100μm for 100X images (i., iii., iv.) and 400μm for 400X images (ii., v., vi.).

## Results and Discussion

We set up crosses between *zpg-cifA;B* males and different female lines *(zpg-cifA;B, zpg-cifA*, and wild type (WT) females), using WT males as control. In all crosses, females mated to *zpg-cifA;B* males showed a striking degree of infertility (only 2–4% viable progeny) compared to controls, suggestive of CI **(Fig. 1b)**. The vast majority of infertile embryos were arrested early in development, while a minority initiated development but did not hatch **(Extended Data Fig. 1).** Embryo cytology revealed the hallmarks of CI^3,4,25^, with most embryos showing early developmental arrest, while others showed fewer nuclear divisions or were arrested later in the blastoderm stage due to mitotic failures **(Fig. 1c)**. Somewhat surprisingly, we did not observe any significant rescue of this infertility when females expressed either *cifA* alone, or both *cifA* and *cifB* **(Fig. 1b).** We also observed a minor (17%) decrease in fertility of *zpg-cifA;B* females compared to their WT and *zpg-cifA* counterparts when mated with WT males **(Fig. 1b)**, strengthening the hypothesis of some reproductive toxicity due to *cifB*.

We speculated that the lack of fertility rescue by *zpg-cifA* could be due to insufficient expression of *cifA* in females, as the rescue effect has been shown to be dose-dependent^26^. To test this possibility, we engineered *cifA* transgenic expression using the *vasa* promoter (**Fig. 1a**), which showed considerably higher expression levels in the germline than the *zpg* promoter (**Fig. 2a**). When mated to *zpg-cifA;B* males, high levels of infertility were observed in both *zpg-cifA* and WT females as above, while fertility was fully restored in crosses with *vasa-cifA* females, demonstrating effective rescue by this transgene **(Fig. 2b).** Combined, these results reveal that CI can be entirely recapitulated in *An. gambiae* mosquitoes by transgenic expression of *cifA* and *cifB*. Incidentally, we also attempted co-injections of *vasa-cifA* and *vasa-cifB* constructs **(Fig. 1a)**, but failed to isolate any *cifB*-expressing progeny, a further confirmation of *cifB’s* possible embryonic toxicity.

**Figure 2:**
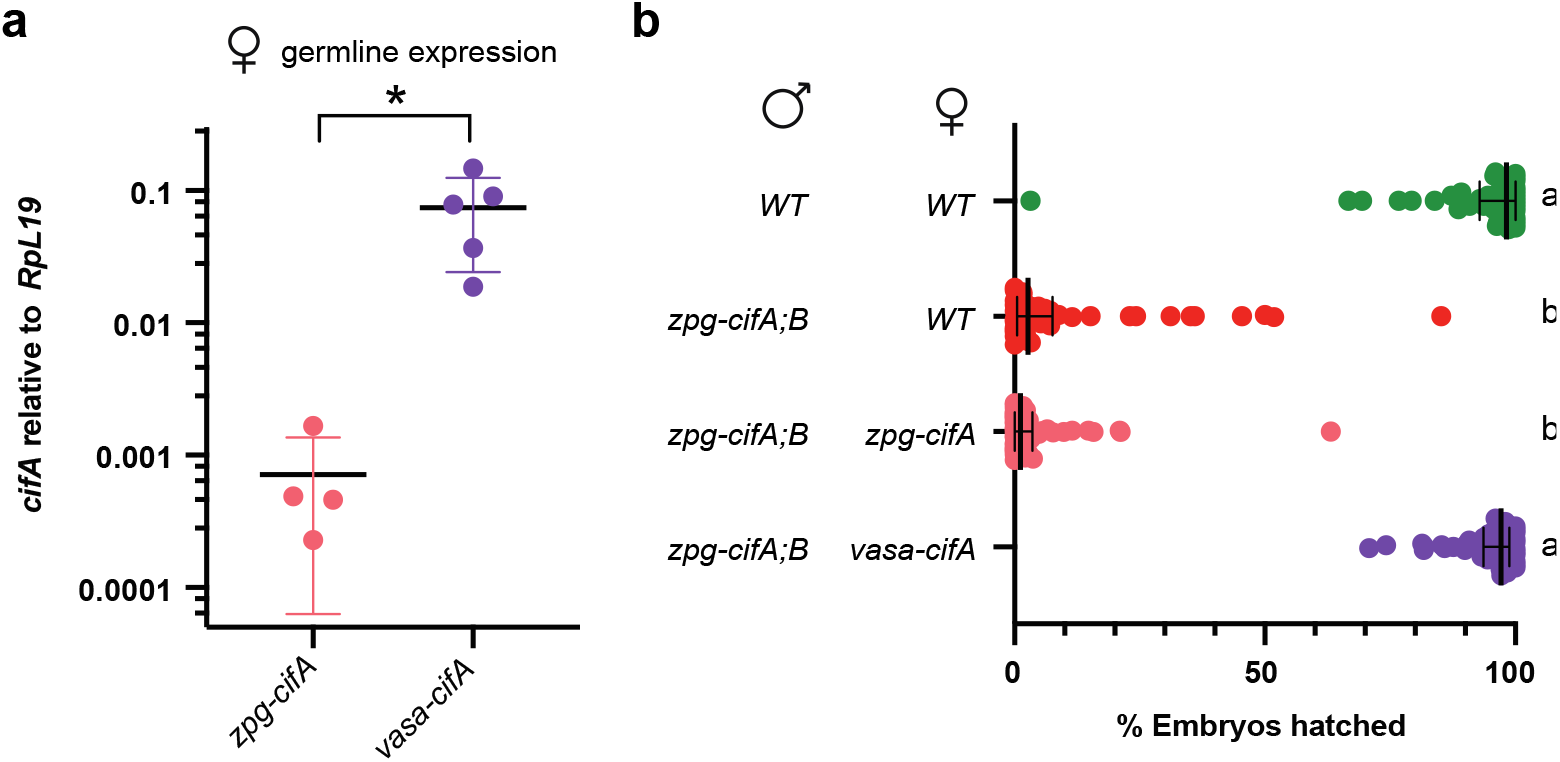
High expression of female *cifA* rescues *cifA;B*-induced CI in *An. gambiae*. (a) Transcript abundance of *cifA* is higher in *vasa-cifA* females compared to *zpg-cifA* females, relative to *RpL*19 (Unpaired t-test (two-tailed), *p*=0.0232, mean and SD are displayed). For each group the *n* is as follows, from left to right: 64, 80. (b) The expression of *vasa-cifA* in females rescues infertility caused by *zpg-cifA;B* expression in males to WT levels, while expression of *zpg-cifA* in females does not (Dunn’s multiple comparison tests, *p*<0.0001 for differences between all statistical groups). Median and interquartile ranges are shown. For each group (top to bottom) the *n* is as follows: 51, 50, 52, 52. Kruskal-Wallis results: H=153.1, p<0.0001, df=3.

Following our observations of potential *cifB* toxicity while establishing the transgenic lines, we investigated whether *cifB* alone is capable of causing CI. Although we could not maintain a *zpg-cifB* colony in the absence of *cifA*, we were able to isolate a limited number of F1 *zpg-cifB* males from natural colony matings between heterozygous individuals. We found that *zpg-cifB* males induced high infertility when mated to WT females, comparable to the infertility levels induced by *zpg-cifA;B* males **(Fig. 3a).** Progeny sired by *zpg-cifA* males were instead fully fertile **(Fig. 3a)**. CI induction did not differ depending on whether *zpg-cifB* males were isolated from *zpg-cifA;B* or *vasa-cifA;zpg-cifB* colonies (denoted *(z)zpg-cifB* or *(v)zpg-cifB*, respectively) **(Fig. 3b).** Once again, *vasa-cifA* expression in females was sufficient to fully rescue sterility caused by *zpg-cifB* males, ruling out CI-independent effects **(Fig. 3b)**. Embryo cytology confirmed the results obtained with *zpg-cifA;B* males, revealing the canonical patterns of CI (**Extended Data Fig. 2**). These findings represent the first conclusive report of conditional sterility induced by *cifB* alone in insects, supporting its independent role as an inducer of CI.

**Figure 3:**
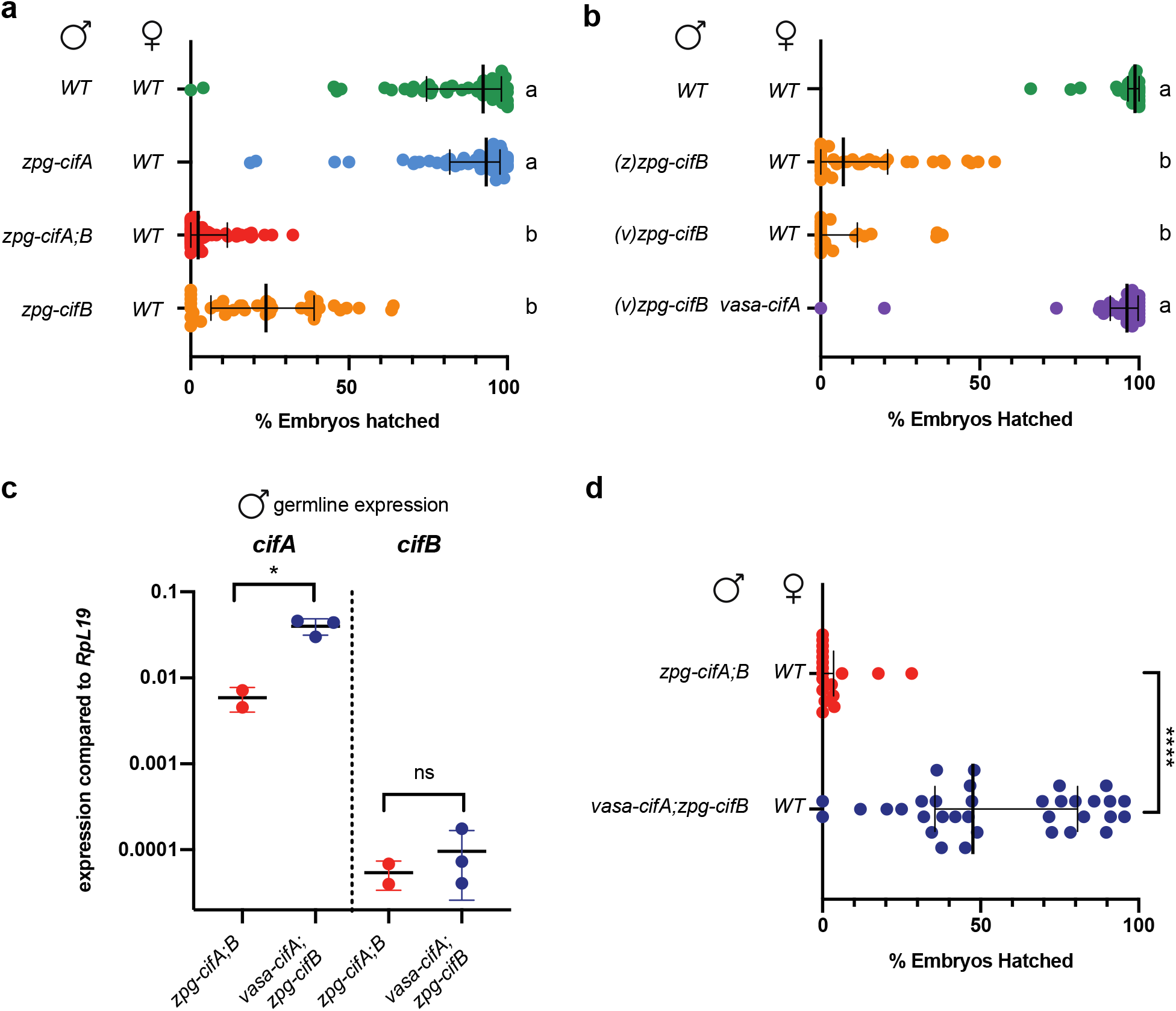
Male *cifB* expression is sufficient to cause CI, while male *cifA* attenuates it. (a) *zpg-cifB* males cause infertility in WT females, while *zpg-cifA* males do not (Dunn’s multiple comparisons, *p*≤0.0001 for differences between all statistical groups). Median and interquartile ranges are shown. For each group (top to bottom) the *n* is as follows: 55, 55, 44, 39. Kruskal-Wallis results: H=133.8, *p*<0.0001, df=3. (b) The expression of *vasa-cifA* in females rescues infertility caused by *(v)zpg-cifB* expression in males, which induce CI to the same extent as *(z)zpg-cifB* males. (Dunn’s multiple comparisons, *p*≤0.0001 for differences between all statistical groups). Median and interquartile ranges are shown. For each group (top to bottom) the *n* is as follows: 36, 39, 24, 32. Kruskal-Wallis results: H=95.08, p<0.0001, df=3. (c) Expression of *cifA* in the male germline is higher in *vasa-cifA* than *zpg-cifA*, (Unpaired t-test (two-tailed), *p*=0.0135, mean and SD are shown), while the expression of *cifB* is similar (Unpaired t-test (two-tailed), *p*=0.4882, mean and SD are displayed). For each group (left to right) the total *n* is as follows: 32, 48, 32, 48. (d) Expression of *vasa-cifA;zpg-cifB* in males causes only partial induction of CI. (Mann-Whitney test (two-tailed), *p*<0.0001. Median and interquartile ranges are shown. For each group (top to bottom) the *n* is as follows: 18, 34.

Given that *vasa-cifA* rescues inviability caused by *cifB* in the embryo while *zpg-cifA* does not, we next asked whether expression levels of *cifA* in males may impact the strength of CI. To this end, we compared fertility of crosses between males expressing either low *(zpg-cifA;B)* or high *(vasa-cifA;zpg-cifB) cifA* levels and WT females **(Fig. 3c).** Intriguingly, *vasa-cifA;zpg-cifB* males were considerably more fertile (median of 48% hatched embryos) compared to *zpg-cifA;B* males (median of 0% hatched embryos) **(Fig. 3d)**, despite similar *cifB* expression levels in these male groups **(Fig. 3c).** These data indicate that high expression of *cifA* in males reduces CI penetrance rather than favoring it, possibly either by limiting CifB activity within the male germline, or by rescuing CifB toxicity in the embryo following transfer of CifA in sperm^27^.

Our finding that *cifB* expression in males is sufficient to induce significant sterility prompted us to investigate toxicity of this factor in females. We designed crosses between WT males and either *zpg-cifA;B* or *vasa-cifA;zpg-cifB* females **(Fig. 4a)** and then characterized egg development and fertility of the *zpg-cifB* F1 female progeny after mating to WT males. We noticed that, in contrast to males (**Fig. 3b**), *cifB*-mediated effects were dependent on the colony of origin. When derived from *zpg-cifA;B* mothers, most F1 *zpg-cifB* females (called *(z^mat^)zpg-cifB)* failed to develop eggs following a blood meal, and only a few females yielded fertile progeny **(Fig. 4b, c).** Additionally, morphological analysis of these ovaries before and after blood feeding showed that follicles were largely absent, suggestive of defects in germline development (**Fig. 4d, e**). On the other hand, when derived from *vasa-cifA;zpg-cifB* mothers, F1 females *((v^mat^)zpg-cifB)* showed intermediate phenotypes, with substantial follicle development, although both fecundity and fertility were reduced compared to WT females **(Fig. 4b-e).** However, when the *cifB* transgene was inherited from *vasa-cifA;zpg-cifB* fathers (**Fig. 4a**), the majority of F1 females *((v^pat^)zpg-cifB)* had ovaries similar to the ones observed in *(z^mat^)zpg-cifB* females, showing remarkably reduced follicle development **(Fig. 4d, e).** As the *zpg-cifB* insertion site and promoter is the same in all these groups, these results reveal rescue effects possibly caused by maternal deposition of CifA from *vasa-cifA*-expressing mothers. *cifB* expression in females is therefore highly deleterious in the female germline when unchecked by the presence of *cifA*, and it appears to act during early stages in germline development based on the capacity for maternally derived *cifA* to rescue these defects.

**Figure 4:**
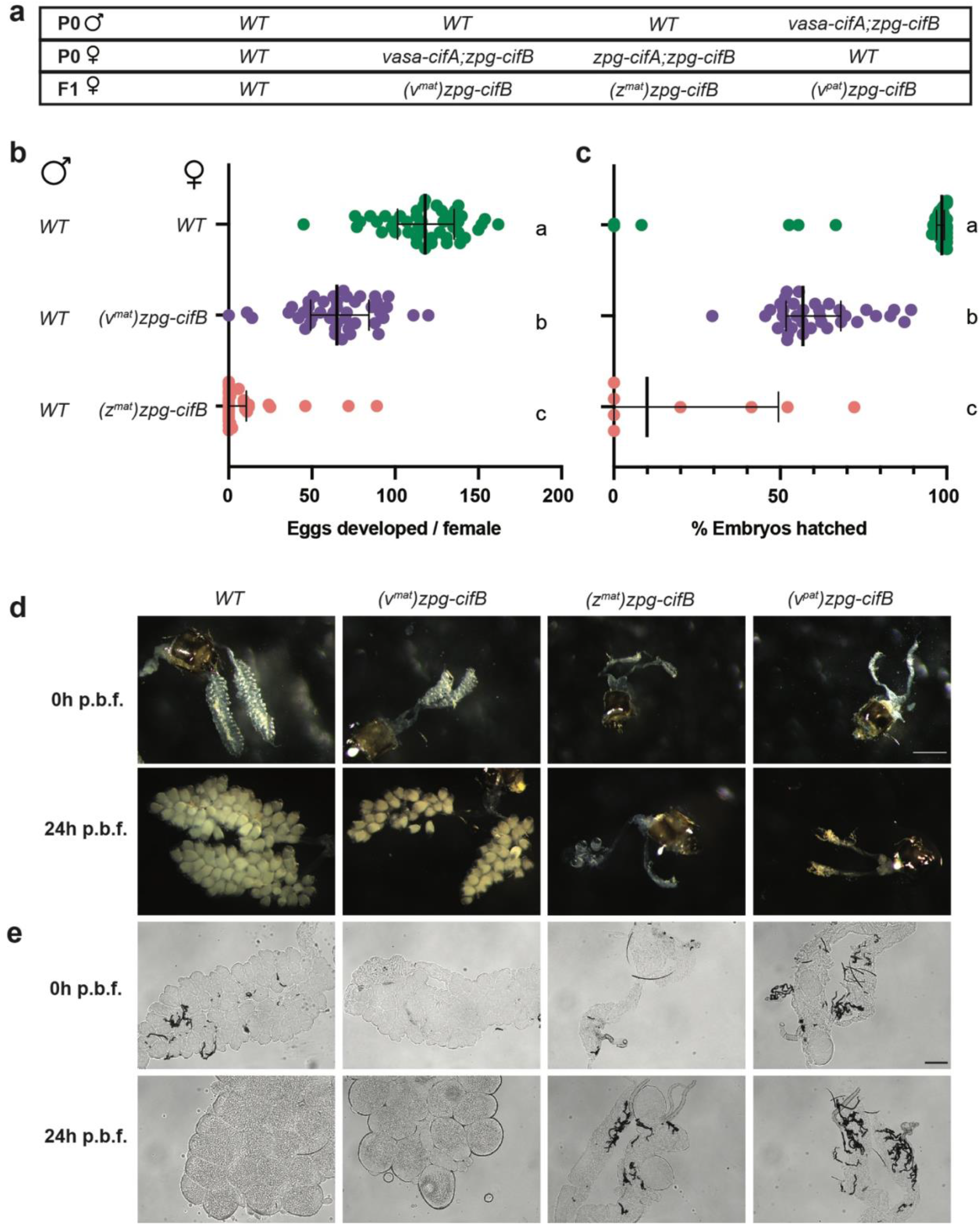
*cifB* expression in females causes severely impaired follicle development in the absence of *cifA*. (a) Crosses were set up to isolate *zpg-cifB* females, F1 progeny derived from either mothers who also expressed *vasa-cifA ((v^mat^)zpg-cifB)* or *zpg-cifA ((z^mat^)zpg-cifB)*, or fathers also expressing *vasa-cifA ((v^pat^)zpg-cifB)*. (b) Egg development is nearly abolished in *(z^mat^)zpg-cifB*-expressing females, while nearly all *(v^mat^)zpg-cifB* females show egg development, although with decreased numbers of eggs compared to WT. (Dunn’s multiple comparisons tests. *p*<0.0032 for differences between all statistical groups). Medians and interquartile ranges are shown. For each group (top to bottom) the *n* is as follows: 23, 20, 24. Kruskal-Wallis test results: H=39.85, p<0.0001, df=2. (c) *(z^mat^)zpg-cifB* and *(v^mat^)zpg-cifB* females show impaired fertility compared to WT females. (Dunn’s multiple comparisons, *p*<0.0001 for differences between all statistical groups). Median and interquartile ranges are shown. For each group (top to bottom) the *n* is as follows: 45, 44, 33. Kruskal-Wallis test: H=91.88, p<0.0001, df=2. (d, e) Ovaries from *cifB* females show severely impaired follicle development unless derived from a *vasa-cifA*-expressing mother, when imaged (d) at either 0h or 24h post blood feeding (p.b.f.) prior to fixing under brightfield microscopy (size marker: 800μm) or (e) after fixing using Differential Interference Contrast (size marker: 100 μm).

Using *cif* genes from *w*Pip in *An. gambiae*, our data show that *cifB* expression is sufficient to induce CI. Previous efforts to generate *cifB^wPip^*-expressing flies were unsuccessful^4^, consistent with our own difficulties of isolating *cifB*-expressing individuals and with our results demonstrating *cifB* toxicity. Our findings are in contrast with results in *D. melanogaster* where both *cifA* and *cifB* from *w*Mel were required to induce CI, and where a *cifB* transgenic line was isolated in the absence of *cifA*^3^. Strain-dependent and/or host-dependent differences may explain the dissimilar findings between CI induction by the Type I *cif* genes in *w*Mel and *w*Pip, but our data nonetheless call into question the universality of the Two-By-One model. In our view, our findings are supportive of a parsimonious toxin-antidote model where CifB is the toxin and CifA is the antidote. Lending support to this, other studies have shown some infertility induced by *cifB* alone in flies, with both *w*Pip’s Type IV *cifB* homolog (also called *cinB)* and *w*Rec’s Type I *cifB* gene, though neither study demonstrated rescue of these effects and thus could not conclude that they were CI related^28,29^. While it is plausible that in some systems CifA may stabilize or even potentiate CifB activity, our results rule out that CifA is always necessary for CI. In fact, *cifA* may even hinder CI induction, as shown by reduced sterility when *cifA* is highly expressed alongside *cifB* in males.

Importantly, we conclusively show that it is possible to both induce and rescue CI in *An. gambiae*, suggesting that it may be feasible to eventually utilize *Wolbachia* to invade these mosquitoes. However, the reproductive toxicity observed in both sexes upon *cifB* expression may partially explain why infections using CI-inducing *Wolbachia* strains have been difficult to establish in laboratory colonies and why these endosymbionts have been detected at low prevalence and intensity in field populations of *Anopheles* mosquitoes^17–22^. Combined with our evidence that high levels of *cifA* are needed to rescue CI in females but attenuate *cifB* activity in males, it emerges that *Wolbachia* may have to fine-tune the relative expression of these two genes in males and females to successfully colonize *Anopheles* mosquitoes. Such a balance undoubtedly would present a hurdle in *Wolbachia*’s invasion of these insect hosts, one that may result in silencing the toxic *cifB* gene through mutation. This hypothesis is supported by the discovery of *cifB* nonsense mutations through sequencing from *w*AnM and *w*AnD strains recently discovered in some *Anopheles* species^30^ as well as by the low prevalence^18,21–23^ and lack of observed CI in *An. gambiae* field populations^23^. With this knowledge, in the future it may be possible to facilitate successful and stable *Wolbachia* colonization of *Anopheles*, for instance by limiting *cifB* toxicity in these mosquitoes via *cifA* germline expression. This would create an avenue to screen *Wolbachia* strains that can block transmission of *Plasmodium* parasites, therefore paving the way to exploit these endosymbionts for malaria control. Finally, the remarkable sterility induced by *cifB* could be utilized for sterile male releases to suppress *Anopheles* populations even in the absence of *Wolbachia* infection, similar to the IIT programs implemented in *Aedes* mosquito control^12,13^. At a time when novel malaria control strategies are urgently needed, our data presents a step towards utilizing *Wolbachia, or Wolbachia*-derived genes, in either population replacement or suppression programs targeting *Anopheles* mosquitoes.

---

**Extended Data Figure 1:**
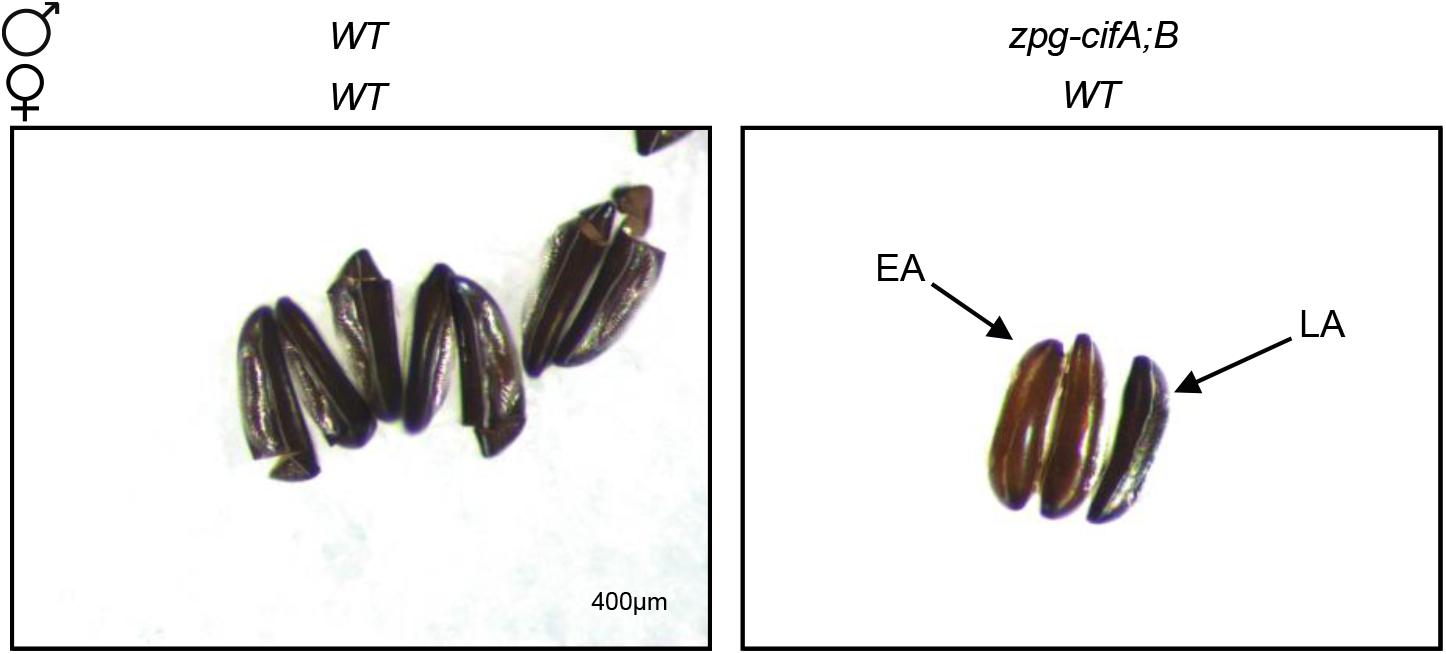
Embryos from *zpg-cifA;B* males show either early or late arrest. Brightfield images of eggs 5 days after oviposition from crosses between WT mosquitoes (left) or *zpg-cifA;B* males and WT females (right). While WT embryos show full development and the standard opening of the hatching cap following larval hatching, embryos from *zpg-cifA;B* males are inviable and arrested either during early development (EA) with a pale brown color, or late development (LA), which show stemmata, but also present severe abnormalities and do not hatch.

**Extended Data Figure 2:**
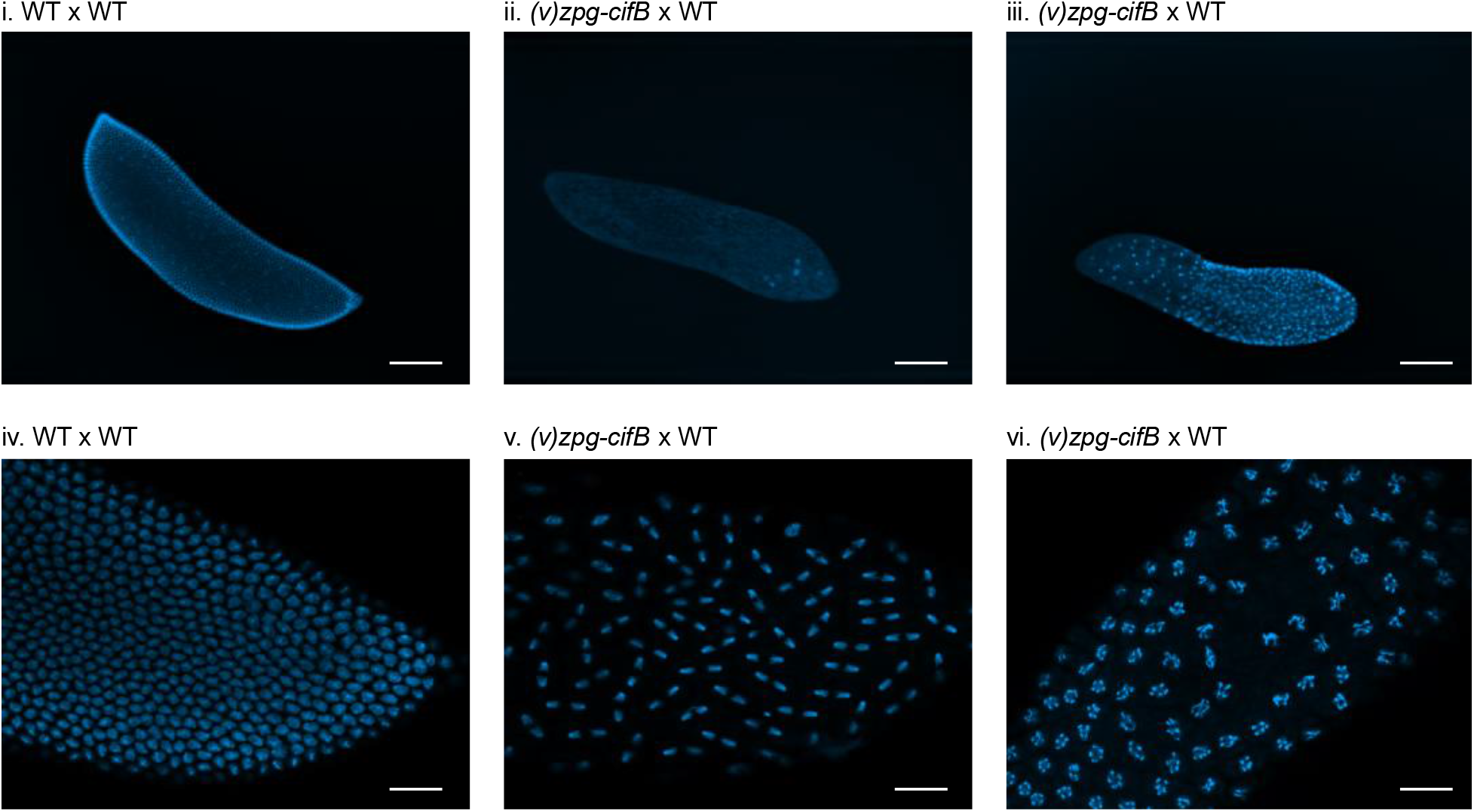
Cytology of the progeny of *cifB* males shows hallmarks of CI. F1 embryos of crosses between either WT or *(v)zpg-cifB* males with WT females were stained for DAPI and imaged 3-4 hours after oviposition. At 100X, while WT controls (i. and iv.) show normal development, *(v)zpg-cifB* embryos show various hallmarks of CI, including (ii.) early arrest, (iii.) regional mitotic failure, and (v., vi.) chromatin bridging or chromosomal abnormalities and delayed or arrested nuclear division. Scale bars represent 100μm for 100X images (i.-iii.) and 400μm for 400X images (iv.-vi.).

## Methods

### Generation of constructs

The amino acid sequences for *cifA* (wPa_0282) and *cifB* (wPa_0283) coding regions from the published *w*PipI Pel strain of *w*Pip from *Culex pipiens*^31^ were codon-optimized by hand for expression in *An. gambiae* using published codon bias information^32^. Gene blocks were ordered from Integrated DNA Technologies (Coralville, IA) using Custom Gene Synthesis to create the desired DNA fragments. Transgenesis constructs were engineered to express the *w*Pip CI genes *cifA* and *cifB* under the control of the germline-specific promoters *zpg (zpg*, AGAP006241) and *vasa (vasa2*, AGAP008578)^33^. The constructs also express a fluorescent marker under control of the ubiquitous *actin* promoter to enable selection of transgenic mosquitoes. Integration into the mosquito genome was mediated by *piggyBac* transposition and rearing lines to homozygosity was accomplished through pupae sorting via fluorescence intensity.

### Embryonic microinjection

*PiggyBac* transgenic construct pairs corresponding to each germline promoter *(zpg-cifA-EYFP* and *zpg-cifB-DsRed*; or *vasa-cifA-EYFP* and *vasa-cifB-DsRed)* were co-injected into the posterior of freshly laid embryos from *An. gambiae* at a concentration of 250ng/μL. Pupae that survived injection were separated according to sex, reared to adulthood, and backcrossed to wild-type G3 to identify and isolate transgenics. A total of 17 EYFP/DsRed double positive F1 transgenics were recovered from the *zpg* promoter-driven CI constructs injections. In contrast, only *vasa-cifA-EYFP* positive transgenics were recovered from the *vasa-cifA/cifB* co-injections. Irrespective of germline promoter, no *cifB* transgenic mosquitoes were identified post-injection.

### Mosquito lines and rearing

*An. gambiae* mosquitoes (species verified by PCR^34^) from the G3 strain and transgenic derivatives of the G3 strain were maintained in a 27°C insectary environment with 70-80% humidity and a 12h light: 12h dark cycle. Adults are given 10% glucose and water *ad libitum* and fed on human blood (Research Blood Components, Boston, MA). Larvae are fed a mixture of Tetramin fish flakes and pellets.

Separate colonies were maintained containing the following transgenes: Colony 1: *zpg-cifA* (Chromosome 3R insertion). Colony 2: *zpg-cifA* (3R); *zpg-cifB* (Chromosome X insertion). Colony 3: *vasa-cifA* (putative Chromosome 2L insertion (2L*)). Colony 4: *vasa-cifA* (2L*); *zpg-cifB* (X). Colony 5: *zpg-cifA* (unknown insertion); *zpg-cifB* (X). To establish Colony 4, *zpg-cifB* males isolated from Colony 2 or 5 were crossed with *vasa-cifA* females from Colony 3. All colonies were maintained as heterozygotes, and screened for fluorescent markers as pupae to select for the presence of transgenes. For all experiments using *zpg-cifA* mosquitoes, Colony 1 was used. For all experiments using *zpg-cifA;B* mosquitoes, Colony 2 was used. For all experiments using *vasa-cifA* mosquitoes, Colony 4 was used. Experiments using *(z)zpg-cifB* mosquitoes used mosquitoes isolated from either Colony 2 or 5. Experiments using *(v)zpg-cifB* males used mosquitoes isolated from Colony 4.

### Crosses and fertility assays

To perform crosses between different transgenic lines, individuals were isolated as pupae from these colonies and their transgenes were identified by their respective fluorescent markers. We did not verify whether individuals were homozygous or heterozygous for their transgenes. Pupae were separated by sex under a dissecting microscope, placed in cages with a ratio between 1:1 and 2:1 males to females, and allowed to eclose in small BugDorm^®^ cages. Natural mating proceeded and mosquitoes were given *ad libitum* access to 10% glucose solution and water for 5–7 days prior to blood-feeding females and allowing oviposition in individual cups lined with filter paper. Once laid, eggs were stimulated daily by spraying water and allowed to hatch for a minimum of 4 days. We then assessed fertility of females by counting and scoring eggs under a Leica M80 dissecting microscope, and additionally noting the presence or absence of hatched larvae. For any female that showed no fertile embryos, we verified her mating status by checking microscopically for the presence of sperm in the spermatheca. For egg development experiments, egg counts for all females were included regardless of whether they had mated or oviposited, while only those that were mated and oviposited were included in fertility experiments.

### RNA extraction and qRT PCR

Male or female reproductive tracts were dissected in pools of 16, collected in TRI reagent (Thermo Fisher Scientific), and stored at −80°C. RNA was extracted according to manufacturing instructions with an additional three ethanol washes of pelleted RNA. Following resuspension, RNA was treated with Turbo DNAse (Thermo Fisher Scientific), quantified with a Nanodrop 2000C (Thermo Fisher Scientific), and then 0.75–2μg were used in a 100μL cDNA synthesis reaction, following standard protocols. We designed primers for qRT-PCR (QuantStudio 6 pro, Thermo Fisher Scientific) using NCBI PrimerBLAST^35^ and after evaluating four different primer sets for *cifB*, we used the following primers for *cifA* and *cifB* at the following concentrations: cifAF: 5’ tcgccgagctgatcgtgaa 3’ (300nM), cifAR: 5’ atcatgtccaggatctccttcttctc 3’ (300nM), cifBF 5’ AGAAGGACCGCCTGATCG 3’ (900nM), cifBR 5’ AGGCTATCGGCGTAGTAGCC 3’ (900nM), RpL19F 5’ CCAACTCGCGACAAAACATTC 3’ (300nM), RpL19R 5’ ACCGGCTTCTTGATGATCAGA 3’ (900nM). Relative quantification was determined using the 2^-(ΔCt)^ equation, using *RpL*19 as the standard. For female *cifA* expression **(Fig. 2a)**, transcript levels were not found to be different between samples from *cifA* only or *cifA* and *cifB* co-expressing individuals, so these data were pooled.

### Microscopy and tissue staining

#### Embryo cytology

Embryos were collected from 10–12 WT females after natural matings with *zpg-cifA;B* males. Four hours after oviposition, embryos were bleached, washed, and dechorionated according to methods by Goltsev *et al*., 2004^36^, and the endochorion was peeled according to methods by Juhn and James, 2012^37^. Embryos were then fixed and stained with DAPI and imaged on a Zeiss Inverted Observer Z1 at 100X or 400X magnification.

#### Brightfield microscopy of embryos

A sample of oviposited embryos were imaged on filter paper at either 5X or 7.5X on a Leica M80 dissecting scope.

#### Brightfield and Differential Interference Contrast imaging of ovaries

Ovaries of 4–7-day-old females were dissected in PBS at either 0h or 24h post-blood-meal and imaged with a Leica M80 dissecting scope at 7.5X magnification for general morphology. After initial imaging, ovaries were fixed in 4% PFA and then mounted in Vectashield^®^ mounting media with DAPI counterstain (Vector Laboratories, Burlingame, CA). Ovaries were then imaged using Differential Interference Contrast on a Zeiss Inverted Observer Z1 at 100X magnification.

### Statistical methods

In all comparisons of fertility or egg development, Anderson-Darling normality tests showed that all data was not normally distributed, so non-parametric Kruskal-Wallis tests with Dunn’s multiple comparisons were used. Distinct samples were used for comparisons. Tests were performed in GraphPad Prism 8. For all fertility or egg development experiments, two to four replicates were performed for all groups. Only one replicate was performed for embryo cytology experiments. For Fig. 4d and 4e, where representative images were selected, one replicate of dissection and imaging of 5–10 individuals from each group was performed. For all qRT-PCR data two to five technical replicates with 16 individuals each were performed for each group, exact *n* given in figure legends.

## Data availability

All data pertaining to this manuscript can be made available upon request.

## Acknowledgements

We would like to thank W. Robert Shaw for careful reading of the manuscript and advice, and Andrea Smidler for help with construct design. This study was funded by a joint Howard Hughes Medical Institute and Bill and Melinda Gates Foundation Faculty Scholars Award to FC (Grant ID: OPP1158190), and by a fellowship of the National Sciences and Engineering and Research Council of Canada to KA. The findings and conclusions within this publication are those of the authors and do not necessarily reflect positions or policies of the HHMI, the BMGF, or the NSERC.

## Author contributions

K.L.A. contributed to literature searches, study design, data collection, data analysis, data interpretation, figure creation, and writing; D.G.A contributed to literature searches, study design, data collection, data analysis, data interpretation and writing, B.C.W contributed to literature searches, study design, data analysis, and data collection, E.K.S. contributed to data analysis and data collection, M.I. contributed to data collection, and F.C. contributed to study design, data analysis, data interpretation, figure creation, writing, and project supervision. The corresponding authors had full access to all of the data in the study and final responsibility in the decision to submit for publication.

